# Executed and imagined grasping movements can be decoded from lower dimensional representation of distributed non-motor brain areas

**DOI:** 10.1101/2022.07.04.498676

**Authors:** Maarten C. Ottenhoff, Maxime Verwoert, Sophocles Goulis, Albert J. Colon, Louis Wagner, Simon Tousseyn, Johannes P. van Dijk, Pieter L. Kubben, Christian Herff

## Abstract

Using brain activity directly as input for assistive tool control can circumvent muscular dysfunction and increase functional independence for physically impaired people. Most invasive motor decoding studies focus on decoding neural signals from the primary motor cortex, which provides a rich but superficial and spatially local signal. Initial non-primary motor cortex decoding endeavors have used distributed recordings to demonstrate decoding of motor activity by grouping electrodes in mesoscale brain regions. While these studies show that there is relevant and decodable movement related information outside the primary motor cortex, these methods are still exclusionary to other mesoscale areas, and do not capture the full informational content of the motor system. In this work, we recorded intracranial EEG of 8 epilepsy patients, including all electrode contacts except those contacts in or adjacent to the central sulcus. We show that executed and imagined movements can be decoded from non-motor areas; combining all non-motor contacts into a lower dimensional representation provides enough information for a Riemannian decoder to reach an area under the curve of 0.83 ± 0.11. Additionally, by training our decoder on executed and testing on imagined movements, we demonstrate that between these two conditions there exists shared distributed information in the beta frequency range. By combining relevant information from all areas into a lower dimensional representation, the decoder was able to achieve high decoding results without information from the primary motor cortex. This representation makes the decoder more robust to perturbations, signal non-stationarities and neural tissue degradation. Our results indicate to look beyond the motor cortex and open up the way towards more robust and more versatile brain-computer interfaces.

## 1 Introduction

Motor neuron diseases, aging-related diseases and accidents can lead to losing a part of or complete muscle control: in the Netherlands alone, 415.000 people are experiencing severe physical disability (2011)[1, 2]. A main predictor of their life satisfaction is their functional independence [3, 4], which could be regained with appropriate assistive tools. An intuitive way to increase functional independence again is to circumvent muscular dysfunction by using brain activity directly as input for control of assistive tools [5, 6]. For a user to achieve sufficient control, the decoder that translates user intent to device motor output needs to have access to rich movement-related neural information. Traditionally, the primary motor cortex has been the primary target, as it has direct downstream output to muscle actuators [7–10]. For example, implantations of microelectrode arrays (MEA) in the hand-knob area of the human primary motor cortex have resulted in state-of-the-art decoders that can decode imagined handwriting at speeds comparable to regular smartphone typing[11].

Electrodes in the primary motor cortex are bringing brain-computer interfaces (BCIs) significantly closer to clinical application, as they provide a rich, spatially local signal from the cortical surface. While the primary motor cortex is widely known for its descending motor neurons and concrete motor commands, there also exist non-direct processes in motor control like feedback processing and attention[12]. Inversely, the descending motor neurons and concrete motor commands do not originate only from the primary motor cortex[13], illustrating that an overly narrow focus on the primary motor cortex in the invasive motor decoding space. It excludes decoding endeavors in a large part of the motor system and fails to take advantage of other potentially informative (sub)cortical areas.

For example, an important area involved in key motor processes are the basal ganglia [14, 15]. The importance of this area is described not only by anatomical and physiological studies, but also by lesions or impairments in this area, caused by diseases such as Parkinsons’ disease. Deep brain stimulation research shows that stimulation in the globus pallidus interna or subthalamic nucleus restores motor function in Parkinsonian patients. From a decoding perspective, it is demonstrated that the signal from the subthalamic nucleus or globus pallidus interna [16] can be used to decode finger clicking, as well as real-time grasp force level decoding from the subthalamic nucleus [17].

Decoding motor processes from non-primary motor areas is explored increasingly more. Recently, several studies demonstrated significant motor decoding results in humans in many different areas, such as the ventral premotor cortex [18], posterior parietal cortex [19–21], somatosensory cortex [18], supramarginal gyrus [18, 20], temporal areas [22], insula [20, 22] and hippocampus [20, 22].

These results indicate that there is decodable movement-related activity in many different areas distributed throughout the brain, however, they decoders only reach performance statistically above chance. It appears as though these areas separately do not provide sufficient information for high-performance decoders. However, all available information might be leveraged by combining activity from multiple areas into a single lower dimensional representation, ultimately increasing the decoders performance [23, 24].

Simultaneously, the decoder could become more robust to perturbations, signal non-stationarities or brain tissue degradation, which is especially important in the target population [25]. This includes people with progressive diseases and/or multiple co-morbidities, meaning that brain regions can become impaired and stop contributing to the decoder. By combining information into a lower dimensional representation, the decoder can compensate for the loss of information from impaired areas and will still have access to sufficient information from other contributing areas.

There are indications that an informative lower dimensional representation of movement related activity exists [24, 26, 27]. Specifically, neural activity might be well represented by a stable general movement related manifold, together with specific sub-manifolds tailored to specific movements [28]. Importantly, this means that there exists a stable covariance structure across movements from distributed recordings, which could be useful for movement decoders.

Therefore, in this work, we expand from the movement decoders based on individual brain areas to a lower dimensional representation of brainwide distributed stereotactic-electroencephalographic (sEEG) recordings, as these electrodes provide high-spatial and temporal resolution throughout the brain [29]. We implemented a Riemannian decoder that directly harnesses the covariance structure of the lower dimensional representation. This way, minimal interference from data transformation or feature synthesis approaches is required. This decoder classifies trials directly based on the distance of the sample covariance matrix per trial to the geometric mean covariance matrix per class (figure 1A). We show significant above chance performance for both executed and imagined movements for nearly all amounts of principal components (figure 2), without the need for areas surrounding the central sulcus. Additionally, we demonstrate that there is similar information in the lower dimensional representation of the lower frequencies between executed and imagined movements (figure 4).

**Fig. 1.**
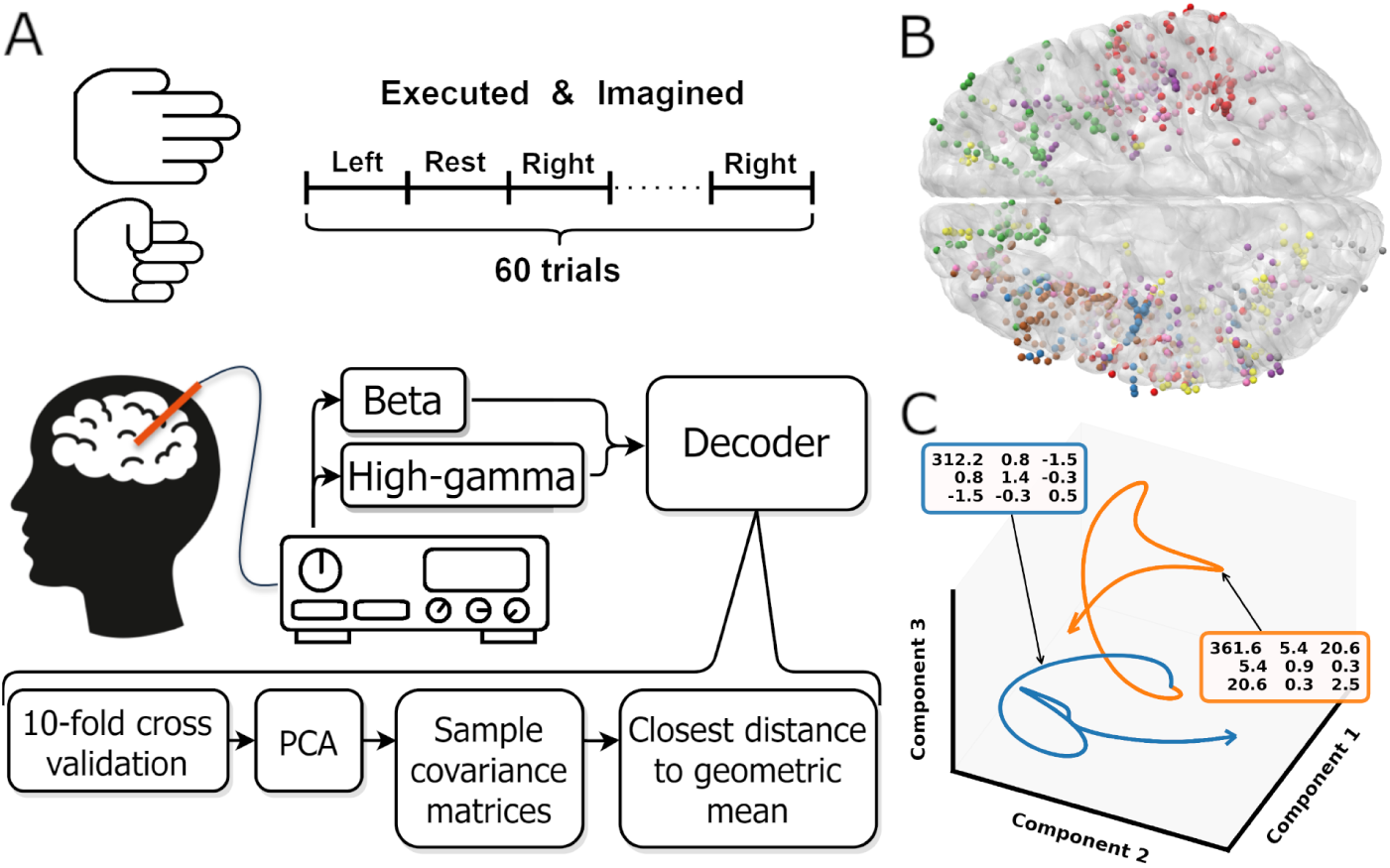
**A)** Overview experimental protocol. **B)** Contact locations of all participants warped onto an average brain. Each color represents contacts from one participant. **C)** Low-dimensional representation of the average movement (blue) and rest (orange) trial. For both trajectories the covariance matrix of the first three components is shown in the colored boxes. The trajectories shown are smoothed by a low pass filter, the unsmoothed trajectories are shown in supplementary figure 1. Note that the trajectories are clearly separated in the space spanned by the first three components.

**Fig. 2.**
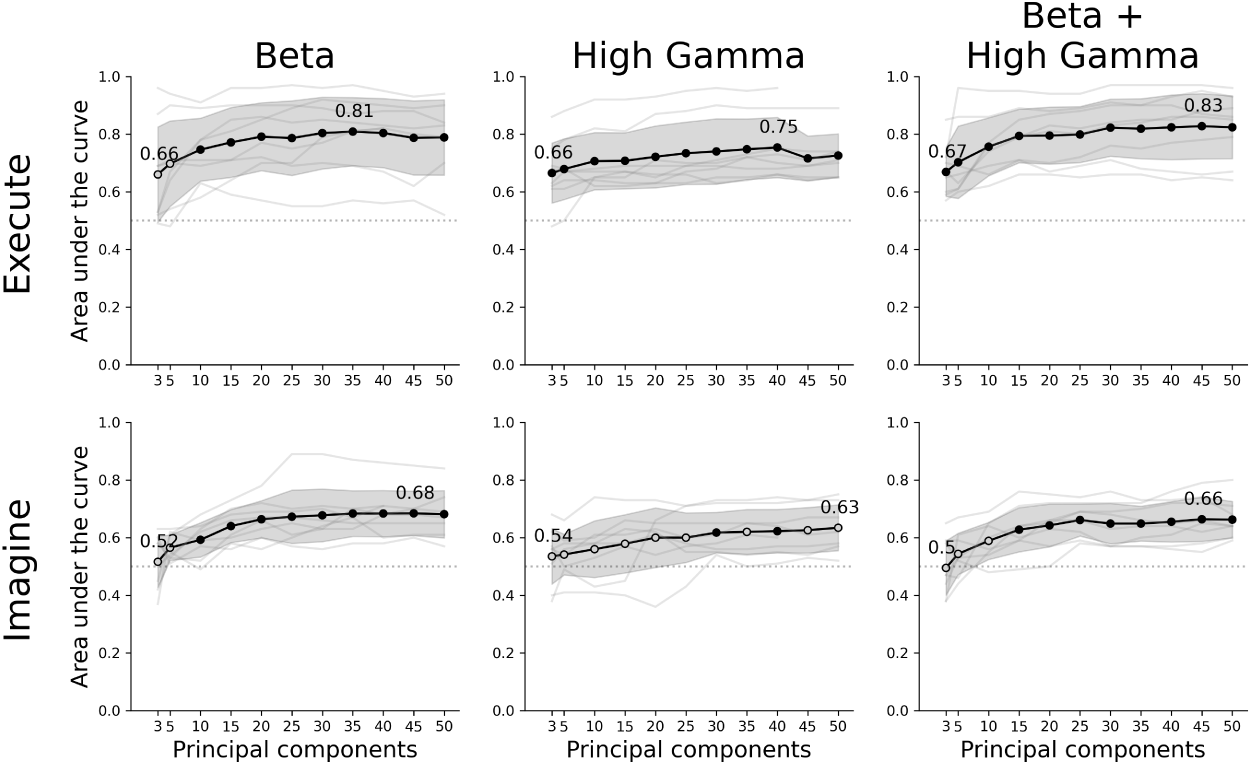
Decoder performance for different movement tasks, frequency features and number of components. The rows show the results of the executed or imagined movement task and the columns each frequency feature set used as input for the decoder. The x-axis depicts the amount of principal components extracted from the data set and the y-axis the AUC score. The light grey lines show the individual average scores over all folds per participant and the black circles are the average scores for a each amount of components. A filled black circle represents an average score that is significantly above chance (corrected for multiple testing), whereas an empty circle is not significant. The grey shaded area shows the standard deviation over participants and the dotted line the chance level (0.5 AUC).

## 2 Results

Eight participants executed and imagined 30 left and 30 right continuous hand grasping movements (figure 1A). We removed all contacts locations surrounding the central sulcus (section 5.1.5), extracted beta and high-gamma frequency components and reduced the data to 3 to 50 principal components. We then trained a Riemannian classifier [30] using 10-fold cross validation on a binary move vs rest task. The model performance was evaluated by the area under the receiver operator characteristic (AUC), and outcomes of statistical tests are corrected for multiple testing.

### 2.1 Executed and imagined movement can be decoded from distributed non-motor brain areas

The classifier decoded executed movements from rest periods significantly above chance for all amount of principal components and frequency features, except beta using 3 or 5 components. The highest performance was achieved by combining beta and high-gamma activity with 45 principal components (0.83 ± 0.11 AUC ± SD, figure 2). Using only beta and high-gamma reached 0.81±0.12 and 0.75±0.10, respectively. As for the imagined movement task, the decoder reached above chance performance for most amounts of components for both beta and beta + high-gamma. However, including only high-gamma produced barely any significant decoding results. Lower amounts of principal components did not reach above chance decoding, specifically: 3 and 5 in beta, 3, 5 or 10 in beta + high-gamma. Overall, performance of decoding imagined movements is lower than executed movements. The maximum performance for imagined movements using beta, high-gamma or beta + high-gamma was 0.68 ± 0.08, 0.63 ± 0.08 and 0.66 ± 0.06, respectively.

### 2.2 A lower dimensional representation is beneficial for decoding from distributed recordings

We trained and tested our Riemannian decoder based on multiple amounts of principal components, ranging from 3 to 50 components. The minimum amount of components that reached above chance decoding was 10 components for all feature sets in executed movement and 15 components in imagined movements (figure 2), except for high-gamma, where only 30 and 40 components reached above chance decoding. In both movement tasks, the decoder performance increased with more components included. From about 15 to 25 components the increase in performance was saturated, though still slowly increasing: the maximum performance was always reached between 35 to 50 components. The informational contribution per electrode to the first principal component, as visualized by the orange and red spheres in figure 3, shows a distributed pattern throughout the brain, illustrating that the sources of the information used by the decoder are not provided by a single area.

**Fig. 3.**
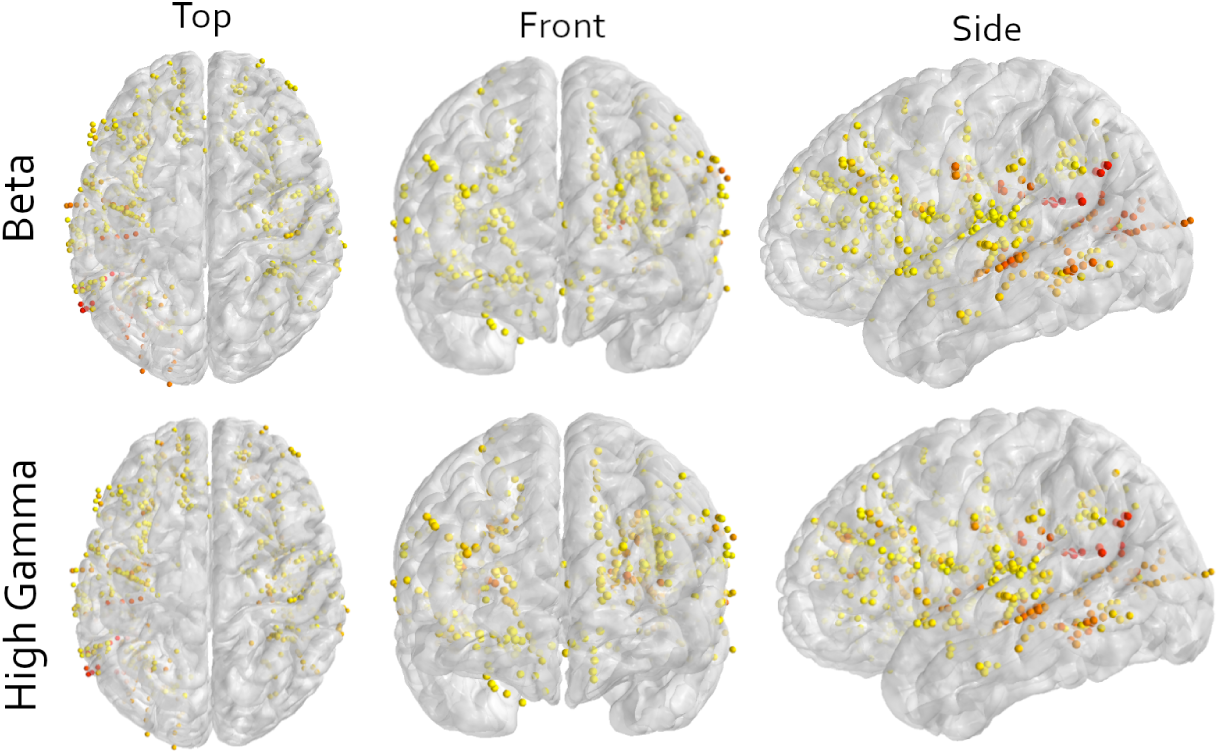
Multi-angle view of all contacts of all participants warped to an average brain for either the beta or high-gamma frequency in the imagined movement task. All contacts in motor cortical areas are excluded (section 5.1.5). Visually, some seem to be located around the central sulcus, which is an artifact most likely caused by warping contacts to an average brain. The color indicates the contribution of that contact to the first principal component, scaled to the explained variance of that component. Yellow means low contribution and red mean high contribution. The image shows that orange and red colors are not bound to a specific area, illustrating the wide distribution of information. Important to note is that a multitude of components is required to achieve the results presented in this work (figure 2, section 2.1), while only the first component is shown in this figure.

### 2.3 Executed and imagined movement share distributed information

If the underlying pattern of executed and imagined movements share information, then the low-dimensional representation should capture similar patterns and achieve above chance decoding. Therefore, we trained the decoder on executed movements, as this signal achieved higher decoding performance, and tested it on imagined movements. For 10 to 30 components in the beta frequency, the decoding performance was significantly above chance. For all other components and in all other frequency bands, the decoder was not able to decode movements above chance (figure 4). This is not surprising, as the decoder was rarely able to decode above chance in imagined movement using only high-gamma, thus indicating that the high-gamma band did not contain much relevant neural information. Further, combining beta and high-gamma resulted in a larger dimensionality without increasing informational content, as indicated by the large increase in standard deviation in beta + high-gamma, thus making it harder for the decoder to fit the data well.

**Fig. 4.**
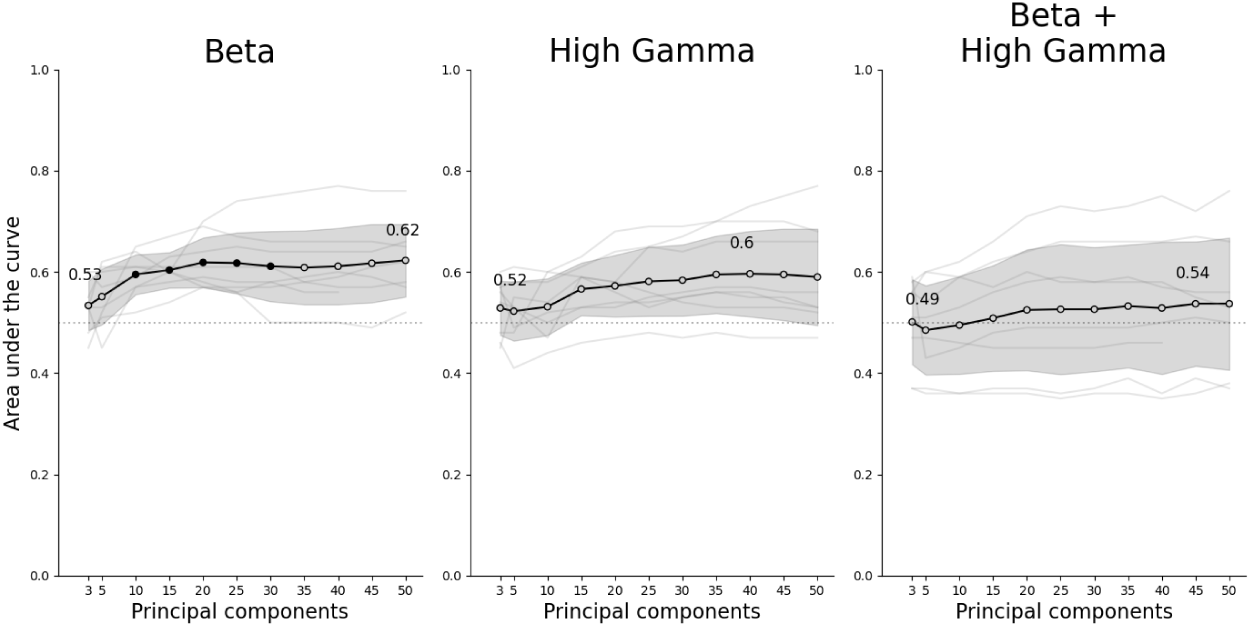
Cross-task results. Decoders are trained on executed movement and tested on imagined movement. Beta yielded above chance decoding for 10 to 30 components, while high-gamma and beta + high-gamma decoding was not above chance.

## 3 Discussion

In this work, eight participants implanted with sEEG electrodes performed executed and imagined movements. We trained a decoder that was able to predict executed and imagined movements based on a lower dimensional representation of distributed non-motor brain areas

Our results contribute to the growing interest in latent motor-related representations in areas other than the primary motor cortex [12]. So far, most studies have either utilized data from local areas using MEAs or ECoG, or used brain-wide recordings using sEEG electrodes but only investigated contributions from grouped contact per mesoscale cortical area [7, 11, 18–20]. Our results extend these investigation by combining the movement-related neural information from all electrodes into a single lower dimensional representation. Decoding from this representation yields good decoding results, specifically in executed movements.

Using a low-dimensional representation has several advantages: 1) The transformation to component space combines the movement-related information from all electrodes into a single representation. While each contact might not contain enough information for good decoding results, combining all these bits of information into a single representation increases the decoders performance. 2) Decreasing dimensionality increases the decoders efficiency, which is essential for translation to a closed-loop system. The amount of calculations for the Riemannian decoder grows exponentially with the amount of dimensions, thus reducing the amount of dimensions drastically improves the computational performance. 3) Combining movement-related information from all contacts makes the decoder more robust to perturbations. Especially in a clinical context, the target population includes many people with progressive brain diseases. This means that the brain tissue could functionally change or deteriorate, potentially causing specific brain regions to decrease or to stop their contribution to the decoder. However, since the representation is a combination of all electrodes, missing one or a few contact is less likely to decrease the decoder performance.

The presented methods utilize any signal that is relevant for the movements performed within the used paradigm, thus no selection is made based on any mechanistic presumption. The relevant signal could also include any other motor related signal, like motor planning, sequencing or decision-making, as well as non-motor information such as attention, stimulus processing, stimulus comprehension or spatial information. While the focus in previous literature is traditionally mostly on motor control, our results indicate that many more information sources could be taken advantage of. However, we consider it unlikely that a single non-motor related process is responsible, since the average amount of used contacts per participants is 95.

We showed that executed and imagined movement share distributed information in lower dimensional space, by training our decoder on executed movements and testing it on imagined movements. This shows that there is similarity between the two tasks and indicates that the underlying processes are also similar. Our results suggest that only the beta frequency is able to capture imagined movements, as it rarely reached above chance decoding in the high-gamma frequency band (figure 2). This is in agreement with a recent brain-wide intracranial speech production study [31], where both beta and high-gamma was informative in overt speech, but only beta activity in imagined speech. Within the beta band, there seems to be an optimum of amount of components that captures the shared information between executed and imagined movements (10 to 30 components, figure 4). Overall, the performance in the cross task decoding is lower. This is not surprising, as we are decoding the shared information, meaning that the decoder utilizes a subset of the information from executed movements. Additionally, the decoder performance in imagined movement was lower when trained and tested on imagined movement in the first analysis. We also identified two technical differences that might influence the results. First, in the cross-task performance, we used executed as training set and imagined as test set, as opposed to the 10-fold cross validation used in the main analysis. Since cross validation generally averages out data set specific noise, this is not the case for a regular train-test setup, which could be a reason for the increased variation in the cross task analysis. Secondly, it is important to note that we did not optimize for performance. In order to investigate the effect of different numbers of components, we chose a decoder that classifies directly on the sample covariance matrix per trial. Through this implementation, we aimed to not further conflate effects on the results with data transformations or feature synthesis required by the decoder. While this provides a better answer to our initial questions, it does mean that if there is relevant high-gamma activity in imagined movements, our decoder pipeline might not be able to capture it.

### 3.1 Limitations

All our participants are diagnosed with refractory epilepsy, a disease of which we do not have a clear picture on how it influences our decoding results. During the monitoring phase in which we perform our measurements, our participants are expected to have as much seizures as possible, albeit no seizures occurred during one of the experimental sessions. After a few days of settling in the monitoring center, medication is reduced and eventually the participants are stimulated in various forms to elicit seizures. Therefore participants often feel drowsy and experience post-ictal discharges. We try to reduce influences as much as possible by visiting as early in their treatment as possible, but we are dependent on the clinical schedule of the patient. Additionally, while we perform our experiments with epilepsy patients, the eventual target population for movement decoders are people with movement impairments. It is unknown if the neural signal in participants that cannot (fully) move anymore is similar enough to decode as well as in epilepsy patients. However, successful studies in paralyzed patients indicates that motor activity can still be decoded [9, 10]. Lastly, we did not objectively control for movements using EMG electrodes during the imagined movement task. This could mean that micromovements or increased muscle tension was present, thus inflating the decoder results.

## 4 Conclusion

In conclusion, we showed that both executed and imagined movements can be decoded from distributed non-motor brain areas using a lower dimensional representation. This shows that there is relevant movement-related information in many areas that might be utilized for decoding purposes, potentially improving BCI applications. Future work should focus on maximizing performance by implementing different decoders and explore more methods to generate a lower dimensional representation. Lastly, future endeavors could transfer to a closed-loop approach to take a further step towards any clinical application.

## Declarations

## 4.1 Acknowledgements

We would like to thank Bart Nolting, Harrie Geeris, Karolina Gasztych, Elly Barten and Stan Hullegie for their invaluable support during recordings.

## 4.2 Author contributions

Conceptualization: MO, CH; Methodology: MO, CH; Software: MO, MV, SG, CH; Validation: MO, PK, CH; Formal Analysis: MO, CH; Investigation: MO, MV, AC, LW, ST, JD, PK, CH; Resources: AC, LW, ST, JD, PK, CH; Data Curation: MO, MV, SG, CH; Writing - original draft preparation: MO, CH; Writing - review and editing: MO, MW, SG, AC, LW, ST, JD, PK, CH; Visualization: MO, MV, SG, CH; Supervision: PK, CH; Project administration: MO, PK, CH; Funding acquisition: PK, CH.

## 4.5 Data and Code availability

Data is publicly available at https://osf.io/xw386, and code on https://github.com/mottenhoff/distributed_motor_decoding

## 5 Methods

### 5.1 Experiment

#### 5.1.1 Participants

Eight participants were included in this work (age 35.8 ± 14.2 years, mean ± SD; 5 male, 3 female, supplementary table 1). All participants are refractory epilepsy patients undergoing presurgical assessment for resection surgery. They were implanted with sEEG electrodes for two to three weeks to monitor seizures and identify the epileptogenic zone. The electrode placement and trajectories were determined solely based on their clinical needs. Participants were implanted with 5 to 14 electrodes containing 42 to 125 recordable contacts.

#### 5.1.2 Ethical approval

The experimental protocol was approved by the institutional review board of Maastricht University and Epilepsy Center Kempenhaeghe (METC 2018-0451). All experiments were in accordance with the local guidelines and regulations and under supervision of experienced healthcare staff. All participants joined the study voluntarily and gave written informed consent.

#### 5.1.3 Protocol

Each participant was asked to continuously open and close their hand for 3 seconds per trial follow by a 3 second rest period. 30 trials were cued per hand, resulting in 60 move and 60 rest trials (figure 1A). The stimuli were presented in random order on a laptop screen that was resting on the participants lap or on a table in front. We ran the protocol for executed and imagined grasping movements.

Participants were instructed to only move their hands and to keep the rest of their body still during executed grasping. For imagined movements, the participants were asked to remain completely still, and the experimenter visually checked if the participants adhered to the instruction. We did not use stricter or more objective methods like electromyography (EMG) to measure any micro-movements or increased muscle tension [32].

In our experience, participants often find it challenging to imagine movements. Therefore, we always preceded the imagined grasping task with the executed grasping task to provide the participant with a fresh memory of the kinematic and proprioceptive sensation of a grasping movement. We assumed it was easier for our participant to recall a mental image of the grasping movement, helping them to perform the imagery task as good as possible. Additionally, the experimenter briefly introduced two potential imagery strategies: kinesthetic or visual [33], but the participants were free to use any strategy that they thought was most effective for them.

#### 5.1.4 Data Recording

Neural activity was recorded by platinum-iridium sEEG electrodes (Microdeep intracerebral electrodes; Dixi Medical, Beçanson, France) using two stacked 64-channel Micromed SD LTM Amplifiers (Micromed S.p.A., Treviso, Italy). The electrodes are 0.8mm in diameter and contain 5 to 18 contacts. The contacts are 2mm in length and have a 1.5mm inter-contact distance and are referenced to a white matter electrode that did not show epileptic activity, visually determined by the epileptologist. All recordings and stimuli were synchronized using LabStreamingLayer[34]. For clarity, throughout this work we refer to ‘electrode’ as the implanted shaft and ‘contact’ for each location on each electrode where activity is measured.

#### 5.1.5 Imaging

The anatomical locations for each contact were determined using the img pipe Python package [35] and parcellation based on the Destrieux atlas [36]. To do so, we coregistered a pre-implantation anatomical T1-weighted MRI scan, parcellated using Freesurfer (https://surfer.nmr.mgh.harvard.edu/), and a post-implantation CT scan. For visualization purposes, the electrodes were warped to average brain from the CVS average-35 atlas in MNI152 space.

To remove motor cortical areas we excluded all contacts of which the determined anatomical label contained the word ‘motor’ or ‘central’ (supplementary data 1). This was a strict exclusion of contacts, meaning that contacts in white matter close to the central sulcus and primary (sensori-)motor cortex are removed as well. Note that the white matter anatomical labels in the Destrieux atlas are based on proximity to labeled grey matter area, introducing some uncertainty of the exact location.

#### 5.1.6 Electrode coverage

In total, 956 contacts on 82 electrodes were implanted in our participants, with electrodes containing a minimum of 5 and a maximum of 18 contacts per electrode. All contacts across participants covered a total of 59 unique grey matter areas with 448 contacts, where the superior insular sulcus is covered the most (*n* = 25) followed by the superior temporal sulcus (*n* = 23) and the middle frontal gyrus (*n* = 23). The remaining contacts are located in white matter (*n* = 408) or unknown areas (*n* = 100). Unknown areas are areas that could not be identified due to various technical reasons. See section 5.1.5 for a technical explanation and supplementary figure 2 for a graphical overview of all areas. Because of a limited number of channels (n=128) that can be recorded by the amplifiers, not all contact could be recorded, reducing the total amount of recorded contacts by 71 (supplementary table 1). The selection of which contacts should be included was made by the epileptologist for clinical reasons. The amount of recorded contacts left after motor and noise removal are shown in supplementary table 1

### 5.2 Decoding

#### 5.2.1 Preprocessing

The data quality of each contact was assessed for excessive noise. First, contacts were flagged if the 50 Hz frequency band power exceeded two times the interquartile range of the signal. Additionally, contacts with a z-scored log square mean value that was significantly higher (*p <* 0.05, assuming normal distribution) than the values in other contacts were flagged for abnormal amplitude (supplementary table 1).

The remaining contacts were detrended, demeaned and band-stop filtered for 50 Hz line noise its and harmonics up to and including 200Hz, using a finite impulse response filter implemented in the MNE python package [37]. Then, we extracted beta (12-30Hz) and high-gamma (55-90Hz) envelope by taking the absolute of the Hilbert transform on the band-passed filtered signal. These frequency bands are chosen as they are known to be movement related and have shown to be effective in decoding studies [7, 38–40]. After preprocessing, the data was split into trials. Left and right hand movement trials were combined into a single movement class.

#### 5.2.2 Decoder

A decoder was trained and tested for [3, 5, 10, …, 50] principle components and beta, high-gamma and beta + high-gamma bands. One participant had less than 50 contacts and could therefore not be evaluated with 50 components. Each component and band combination was trained and evaluated as follows: first, the data was split using 10-fold cross validation. On the training data, the data was standardized over all included trials per fold and a principal component analysis was performed. The learned transformation was subsequently used to transform the training and test fold to the specific amount of principal components. For the cross task performance, the decoder used all data per participant from the executed movement task as training set and all data from the imagined movement task as test data.

After transformation into the components space, the sample covariance matrix for each trial was calculated and regularized by the ledoit-wolf lemma [41]. Then, the geometric mean per class was calculated based on the Kullback-Leibler divergence. Trials were then classified by selecting the class with the shortest distance to class geometric mean. For the calculations, we used the pyRiemann implementation [42].

We choose a Riemannian approach as these decoders have shown promising results by using straightforward methods in surface EEG [30, 43], though no application has been published using sEEG thus far. Additionally, because Riemannian decoders classify directly on the sample covariance matrix, it does not require any further transformations or feature synthesis. As motor behavior should be captured by the covariance in the neural recordings, we hypothesized that these decoders would also perform well on our invasive data.

#### 5.2.3 Evaluation

We evaluated the decoder by the area under the receiver operator characteristics (AUC). For both the main decoder and cross task results, we tested statistical significance against chance level (mean *AUC* = 0.5) using a one sample t-test and corrected for multiple testing using Bonferroni correction. For the control analysis for motor cortical areas, we used a Wilcoxon signed rank-test (bonferroni corrected, *n* = 66) to compare the difference in performance with and without motor cortical areas.

