## Supplementary data for "Executed and imagined grasping movements can be decoded from lower dimensional representation of distributed non-motor brain areas"

| # | Age | Sex | Sample rate | Electrodes | Contacts | | Noise | | Motor |
|---|---|---|---|---|---|---|---|---|---|
| | | | | | Executed | Imagined | Executed | Imagined | |
| 1 | 16 | M | 2048 | 14 | 116 | 116 | 8 | 8 | 10 |
| 2 | 47 | M | 1024 | 11 | 110 | 108 | 20 | 22 | 0 |
| 3 | 52 | M | 1024 | 6 | 54 | 52 | 65 | 67 | 9 |
| 4 | 22 | F | 1024 | 5 | 42 | 44 | 85 | 83 | 0 |
| 5 | 20 | F | 1024 | 11 | 106 | 105 | 13 | 14 | 6 |
| 6 | 40 | M | 1024 | 12 | 117 | 117 | 13 | 13 | 0 |
| 7 | 55 | F | 1024 | 12 | 108 | 105 | 15 | 18 | 8 |
| 8 | 34 | M | 1024 | 11 | 108 | 106 | 17 | 19 | 3 |

**Table 1**: Participants and their electrode configurations. 'Contacts' denotes the amount of contacts after noise and motor removal. 'Motor' denotes the amount of contacts located in an area surrounding the central sulcus.

### Data 1: List of removed areas surrounding the central sulcus

- ctx-rh-paracentral
- ctx-rh-precentral
- ctx-rh-postcentral
- wm-lh-paracentral
- wm-lh-postcentral
- wm-lh-precentral
- wm-rh-paracentral
- wm-rh-postcentral
- wm-rh-precentral
- ctx-lh-G_paracentral
- ctx-lh-G_postcentral
- ctx-lh-G_precentral
- ctx-lh-G_subcentralctx-lh-S_central
- ctx-lh-S_paracentral
- ctx-lh-S_postcentral
- ctx-lh-S_precentral-Inferior-part
- ctx-lh-S_precentral-Superior-part
- ctx-lh-S_subcentral_ant
- ctx-lh-S_subcentral_post
- ctx-rh-G_paracentral
- ctx-rh-G_postcentral
- ctx-rh-G_precentral
- ctx-rh-G_subcentralctx-rh-S_central
- ctx-rh-S_paracentral
- ctx-rh-S_postcentral
- ctx-rh-S_precentral-Inferior-part
- ctx-rh-S_precentral-Superior-part
- ctx-rh-S_subcentral_ant
- ctx-rh-S_subcentral_post
- wm-lh-G_paracentral
- wm-lh-G_postcentral
- wm-lh-G_precentral
- wm-lh-G_subcentral
- wm-lh-S_central
- wm-lh-S_paracentral
- wm-lh-S_postcentral
- wm-lh-S_precentral-Inferior-part
- wm-lh-S_precentral-Superior-part
- wm-lh-S_subcentral_ant
- wm-lh-S_subcentral_post
- wm-rh-G_postcentral
- wm-rh-G_precentral
- wm-rh-G_subcentral
- wm-rh-S_central
- wm-rh-S_paracentral
- wm-rh-S_postcentral
- wm-rh-S_precentral-Inferior-part
- wm-rh-S_precentral-Superior-part
- wm-rh-S_subcentral_ant
- wm-rh-S_subcentral_post
- ctx_lh_G_and_S_paracentral
- ctx_lh_G_and_S_subcentral
- ctx_lh_G_postcentral
- ctx_lh_G_precentral
- ctx_lh_S_central
- ctx_lh_S_postcentral
- ctx_lh_S_precentral-inf-part
- ctx_lh_S_precentral-sup-part
- ctx_rh_G_and_S_paracentral
- ctx_rh_G_and_S_subcentral
- ctx_rh_G_postcentral
- ctx_rh_G_precentral
- ctx_rh_S_central
- ctx_rh_S_postcentral
- ctx_rh_S_precentral-inf-part
- ctx_rh_S_precentral-sup-part
- wm_lh_G_and_S_paracentral
- wm_lh_G_and_S_subcentral
- wm_lh_G_postcentral
- wm_lh_G_precentral
- wm_lh_S_central
- wm_lh_S_postcentral
- wm_lh_S_precentral-inf-part
- wm_lh_S_precentral-sup-part
- wm_rh_G_and_S_paracentral
- wm_rh_G_and_S_subcentral
- wm_rh_G_postcentral
- wm_rh_G_precentral
- wm_rh_S_central
- wm_rh_S_postcentral
- wm_rh_S_precentral-inf-part
- wm_rh_S_precentral-sup-part
- ctx-lh-primary-motor
- ctx-lh-premotor
- ctx-rh-primary-motor
- ctx-rh-premotor

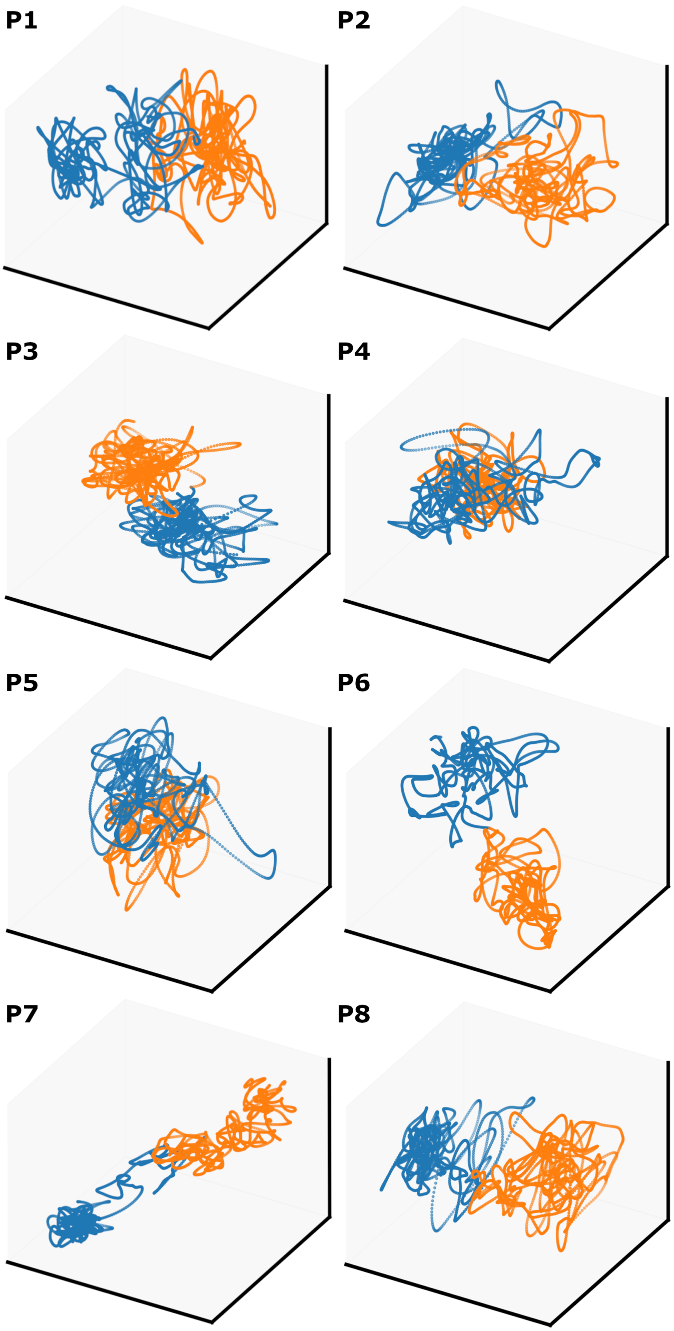


**Fig. 1**: Unsmoothed trajectories in components space. Calculated from beta activity in executed movements. The trajectories of P8 are the same as in Figure 1c.

| | 3 | 5 | 10 | 15 | 20 | 25 | 30 | 35 | 40 | 45 | 50 |
|---|---|---|---|---|---|---|---|---|---|---|---|
| Exec - Beta | 0.98 | 0.71 | 0.83 | 0.89 | 0.84 | 0.90 | 0.98 | 0.98 | 1.00 | 0.95 | 0.94 |
| Exec - High-gamma | 0.86 | 0.79 | 0.93 | 0.86 | 0.95 | 0.75 | 0.91 | 0.89 | 0.92 | 1.00 | 0.97 |
| Exec - Beta + High-gamma | 0.83 | 0.73 | 0.87 | 0.76 | 0.82 | 0.75 | 0.63 | 0.91 | 0.91 | 0.30 | 0.72 |
| Imag - Beta | 0.65 | 0.47 | 0.44 | 0.88 | 0.74 | 0.84 | 0.84 | 0.66 | 0.87 | 0.78 | 0.85 |
| Imag - High-gamma | 0.80 | 0.97 | 0.75 | 0.77 | 0.98 | 0.92 | 0.93 | 0.82 | 0.79 | 0.80 | 0.76 |
| Imag - Beta + High-gamma | 0.85 | 0.78 | 0.79 | 0.75 | 0.80 | 0.79 | 0.75 | 0.75 | 0.84 | 0.83 | 0.88 |

**Table 2**: P-values for significance test between the performance using all electrodes and the performance without motor cortical areas. None of the tests were significant, meaning that the null hypothesis that both groups come from the same distribution cannot be rejected.

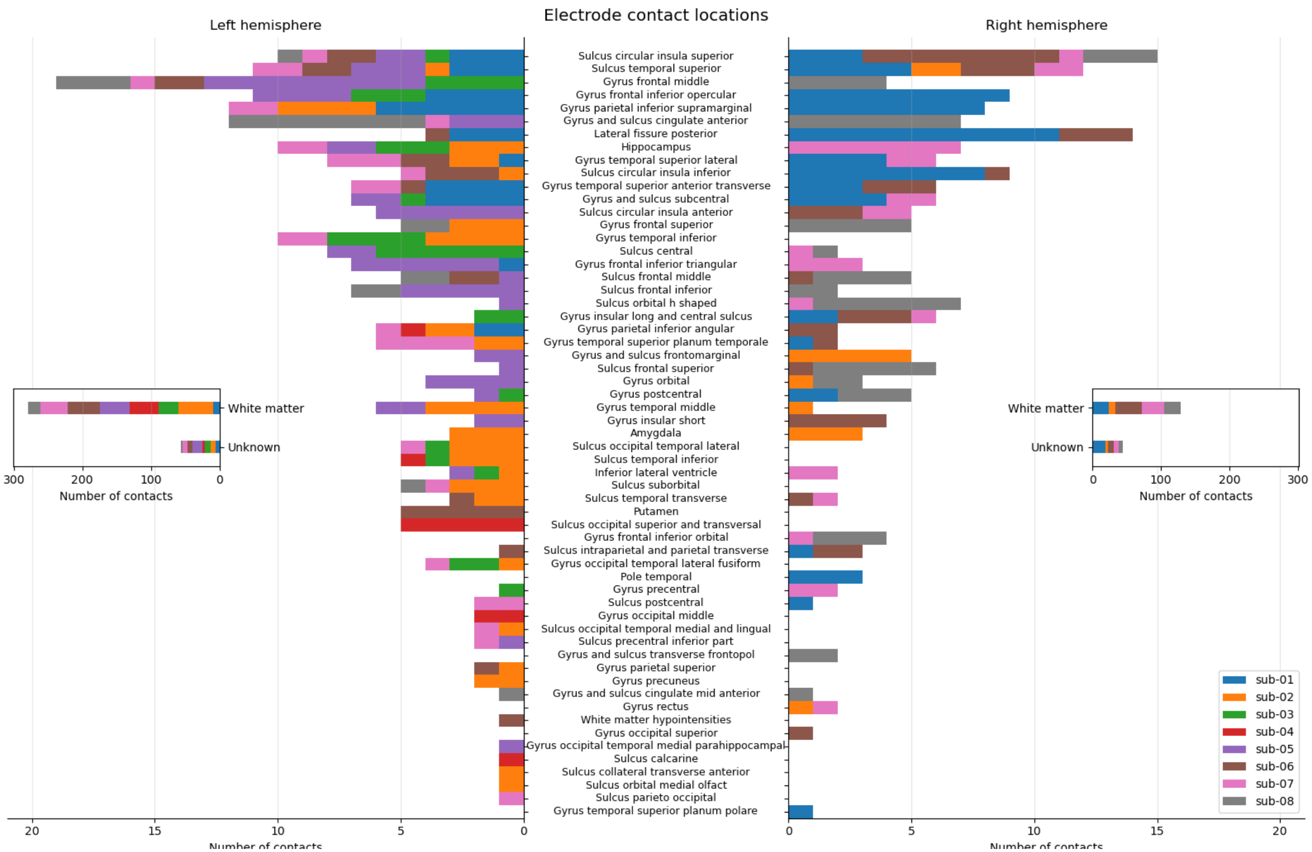


**Fig. 2**: All areas in the brain captured per contact. Note the large X-axis on the inset, most contacts are either in white matter or could not be identified.
